# Late-Talking Children Talk More? A Machine Learning Approach to Speech Act Analysis in Early Childhood

**DOI:** 10.1101/2025.09.30.679667

**Authors:** Girwan Dhakal, Hongsheng He, Sharlene Newman, Yanyu Xiong

## Abstract

Speech acts shape early language development and social cognition, yet little is known about how late-talking (LT) children use them to achieve communicative goals. We compared LT and typically developing (TD) preschoolers (1;09–6;00) across nine dyadic English corpora, using a Conditional Random Field model to annotate speech acts. We analyzed speech act distributions, hierarchical relations, and contingent responses to assess production and comprehension. TD children produced more declarative statements and wh-questions, whereas LT children produced more unclear word-like utterances and showed reduced comprehension ability (1;09–2;07). Speech acts classified LT and TD groups with 72.3% accuracy, improving to 76.6% with linguistic and demographic features. Classification was driven by co-occurring patterns of speech act frequencies. LT children showed delayed onset of speech acts but employed more speech acts after 3;09, focusing on speaker-centered goals, whereas TD children favored collaborative use of speech acts, revealing complex dynamics in the development of communicative skills.

---

A speech act refers to a goal of uttering a proposition by a speaker, including the production of an utterance (locutionary act), the speaker’s intent in producing the utterance (illocutionary act) and the effect of the utterance on the hearer (perlocutionary act) (Levinson, 1983; Searle, 1976; Searle & Vanderveken, 1985; Cameron-Faulkner 2014). It is tied to pragmatic functions of communication, such as declaration, request, response and assertion, etc. Speech acts are pivotal for both child language development and overall developmental outcomes later in life (Ninio et al., 1994) due to children’s innate predispositions to interact with social environment. For typically developing (TD) children, the onset of speech acts occurs at a much earlier stage compared with other linguistic skills during infancy and undergoes continuous development even after language skills have matured (Stephens & Matthews 2014). Before 9 months, infants mainly rely on proto-imperative and proto-declarative communication in prelinguistic dyadic interaction (Bates et al., 1975; Bates, 1976; Bruner, 1975) which gradually develops into triadic interaction with joint attention and mutual understanding of intentions (Tomasello, 2003, 2008; Carpenter, 2012, Bakeman & Adamson, 1984).

For the children with delayed expressive language development, how speech acts are affected remains far less known. Prior studies reported that late talking (LT) children before age 3;00 produced fewer speech acts to convey communicative intents and maintain interactions than the age-matched typically developing (TD) children (Rescorla & Merrin, 1998; Paul & Shiffer, 1991), but the differences decreased afterwards (MacRoy-Higgins & Kliment, 2017). The other cohort of studies focused on nonverbal communication like gestures and word-gesture integration of the LT children between ages 1;04–3;00. They found that the children with persistent delay deficit did not rely on compensatory usage of gestures as late bloomers after 3;00 (Thal et al., 1991; Thal & Tobias, 1994). However, because the frameworks for defining and categorizing speech acts vary across these studies (e.g. rate of communication/joint attention, Dore’s Primitive Speech Acts, conversation management and gesture-based communication acts), the findings are difficult to compare directly as they tap different subsets of pragmatic skills. The absence of a comprehensive speech act scheme that can be consistently applied prevents researchers from forming a coherent understanding of the characteristics of delayed speech act development.

The Inventory of Communicative Acts and its abridged version (INCA-A) (Ninio & Wheeler, 1984) is a coding system of structured discourse that incorporates theoretical Speech Act Theory (Searle, 1976; Searle & Vanderveken, 1985), findings of interactive studies (Goffman, 1961) and caregiver feedback. It has been applied to TD children and caregiver-child speech analysis (Snow et al., 1996; Trautman & Rollins, 2006; Zhao et al., 2024). This system codes 67 communicative acts at both speech act and interchange levels, such as confirmation and clarification, etc., that captures paired acts in conversational routines, is robust and sensitive to the developmental pattern of acts in the TD children, the communicative functions of both caregivers and children and cultural differences. Recent research by Nikolaus et al. (2022) tested the feasibility of using language models to conduct automated INCA-A labeling in large-scale CHILDES datasets. Their models were tested on a small manually coded dataset (Snow et al., 1996) and validated on the large CHILDES corpora. The best performing model— Conditional Random Field (CRF), achieved accuracy of up to 72.33%, showing good accuracy in automatic labeling. Bergey et al. (2024) used the trained CRF model to label 727 caregiver-child interaction sessions of the Language Development Project corpus. They found that the repertoire of communicative acts in the TD not only increased in children from 1;02 to 4;10 but also showed diverse individual differences.

In contrast, the empirical literature on automated INCA labeling to analyze the interactive speech of the LT children is rather limited and under-documented. Utilizing the increased number of open access LT datasets, our current study seeks to address four core research questions in the field, using large language models and machine learning techniques.

First, what are the robust differences in speech acts between the preschool LT and TD children in terms of the speech act type (illocutionary force) and their conversational function across a variety of adult-child interactions? By applying the automatic labeling model trained on large corpora, we aim to capture a broader spectrum of pragmatic functions (up to 55 speech act categories) than the older speech act schemes (8-12 categories) and understand the acts from two dimensions – speech intention and its functional role in an interaction sequel, which are not distinguished in earlier studies (Paul & Shiffer, 1991; MacRoy-Higgins & Kliment, 2017; Rescorla & Merrin, 1998). The approach can help us to derive robust and reproducible classification of communicative acts.

Second, what are the predictive speech acts that can distinguish the LT from TD children? Using the classification techniques of machine learning, we are interested in knowing what speech acts can effectively separate the LT from TD children in the unseen data after cross-validations. Going beyond merely describing the different usage rates of speech acts between the two groups, our study pursued to identify reliable speech act markers that may flag early and targeted intervention for at-risk children before school entry.

Third, what are the differences in speech act emergence between the LT and TD children in terms of production and comprehension? Previous studies have shown that speech acts do not develop in parallel in production and comprehension for the TD children (Snow et al., 1996; Nikolaus et al., 2022). The overall precedence of understanding more speech acts over production indicates that well-established receptive knowledge of functional roles of various speech acts may influence the full development of pragmatic skills in production at a later stage. For the LT children, evidence seems to converge that children with both receptive and expressive delays are at a higher risk of developmental language disorder (DLD) compared to those with their comprehension skills within a normal range (Marchman & Fernald, 2013; Nouraey et al., 2021; Roos & Weismer, 2008). However, little is known of whether the LT children will show different age of emergence of speech acts from the TD children.

Fourth, evaluating the specificity and sensitivity of the INCA to label the speech acts of the LT children. Nikolaus et al. (2022) reported their model performance of classifying different speech acts in the INCA coding scheme for the TD children was affected by either ambiguous definition of some categories or the hierarchical relations among some speech acts. They observed improvement of model performance in their exploratory analysis by collapsing certain overlapping categories. Our study used the data-driven approach to assess the ranked importance and internal relationships among the speech acts to discriminate the LT from TD children. The aim will help us understand the extent to which specific speech acts are differentially important in characterizing the LT versus TD children.

## Methodology

### Data Source

We analyzed nine publicly available English corpora of children with language impairment issues in the CHILDES repository. The longitudinal datasets included Ellis Weismer (Weismer et al., 2013), Hargrove (Hargrove et al., 1986), Rescorla (1989), UCSD (Ziegler et al., 1990), Conti Ramsden 1 and Conti Ramsden 2 (Conti-Ramsden & Dykins, 1991). The cross-sectional datasets included ENNI (Schneider et al., 2006), Eisenberg Guo & Guo (2013) and Bliss (1988). To control unequal weighting in longitudinal datasets, we stratified ages into four bins (1;09–2;07, 2;08–3;09, 3;09–4;10, 4;11–6;00) and restricted each child to a single age bin selected at random from those in which they appeared. Without this adjustment, children with many transcripts within an age bin would dominate aggregate speech act distributions, biasing LT–TD comparisons. Within the selected bin, if a child had multiple transcripts, we computed speech act proportions per transcript and then averaged them. This two-step approach helps mitigate potential biases arising from differences in transcript length, interaction context, or the adult interlocutor. By assigning equal weight to each transcript during averaging, we reduce the influence of any single session dominating a child’s overall speech profile. This ensures unbiased contribution per child, preserves developmental structure, and allows us to analyze age-related trends in speech act usage. It also enables both the replication of key findings from Nikolaus et al. (2022) and the exploration of speech act differences between typical developing and late-talking children across developmental stages. Children with only one transcript in a longitudinal corpus were treated as cross-sectional cases and contributed that single transcript. After sampling the longitudinal datasets, there were 126 samples for ages 1;09–2;07, 32 samples for ages 2;08–3;08, 47 samples for ages 3;09–4;10, and 60 samples for ages 4;11–6;00.

The datasets used in our analysis span both longitudinal and cross-sectional designs, covering a range of ages, activities, and language development profiles. The Ellis Weismer dataset (Ellis Weismer et al., 2013) is a 5-year longitudinal study of 112 children (56 late talkers and 56 typically developing) captured during toy play between ages 2;06 and 5;06, with a mean sample age of 3;00. Bliss (Bliss, 1988) is a cross-sectional corpus comparing children with specific language impairment (SLI) to typically developing peers, consisting of 10 transcripts from children with mean age 4;02. The Eisenberg-Guo dataset (Eisenberg & Guo, 2013), based on picture description tasks, includes 34 preschool-aged children from suburban New Jersey, with a mean age of 3;06. ENNI (Guo & Schneider, 2016) comprises 126 transcripts from children aged 4;00–9;00(mean age 5;01) participating in narrative storytelling; 100 are typically developing and 26 have language impairments.

Several additional datasets focus specifically on late talkers and language-impaired children through naturalistic play. Hargrove (Hargrove et al., 1986) includes longitudinal transcripts from five late-talking male children aged 3;0–6;0 (mean age 3;04). Rescorla (Rescorla, 1989) offers cross-sectional toy play samples from 65 children (mean age 4;04), including 39 late talkers, with a total of 145 transcripts. UCSD-SLI is another longitudinal corpus of 15 late-talking children (10 males, 5 females) with a mean age of 4;07. Conti-Ramsden 1 (Conti-Ramsden & Dykins, 1991) includes 43 language-impaired children matched with younger siblings on Mean Length of Utterance (mean age 3;01), while Conti-Ramsden 2 captures two years of play-based interaction between three children with SLI and their typically developing siblings, with a mean age of 4;08. Across these corpora, toy-play and picture-based activities were the dominant elicitation contexts, supporting consistent analysis of naturalistic language production.

The chosen datasets offer a broad age range and their diagnosis for speech impairment is clinically verified. Each transcript contains manually transcribed child utterances. For the analysis, we only considered English speaking children who are 6;00 or younger. We used the childespy API (Mankewitz et al., 2020) to programmatically retrieve utterance data from each dataset, identified the control and impaired groups within each dataset and assigned a label to each utterance. Finally, all the datasets were combined into a table for analysis.

**Table 1.**
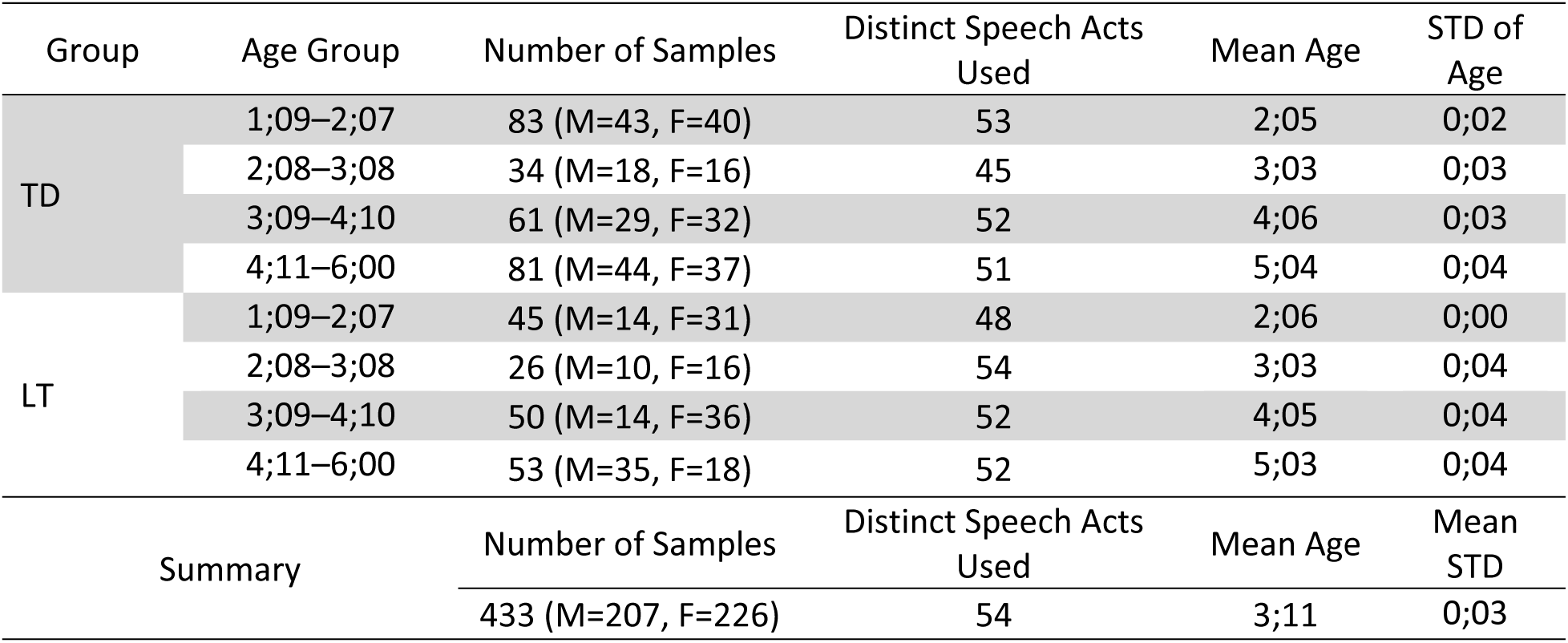
Dataset Summary.

### Speech Act Labeling

To create the working definitions of speech acts, we incorporated the idea of a hierarchical speech act system (Zhao et al., 2024) by proposing three-tier coding taxonomy to further simplify and organize the INCA-A scheme (Appendix Table 1). The top tier includes 7 broad categories (interaction regulation, social rituals, joint attention, information exchange, discourse organization and miscellaneous), and each category encompasses illocutionary acts (*i.e.* requests, responses and asserting, etc.) as the second tier. Each illocutionary act can be realized by different types of speech acts at the third tier depending on the context of interaction. We placed speech acts that have multiple functions into the categories they can be associated with. For example, ST (make a declarative statement) is grouped under both ‘joint attention’ and ‘information exchange’ as it is frequently used in interaction for commenting a co-reference and/or making assertion in responses.

We used the pretrained conditional random field (CRF) model from Nikolaus et al. (Nikolaus, 2024) to automatically annotate children’s utterances with INCA-A speech act labels. A CRF model is a probabilistic model often used to label a sequence of inputs. The model considers the local features such as the utterances, parts-of-speech tags and so on but also contextual dependencies. (Sutton & McCallum, 2010) This makes the model appropriate for speech act annotation. The CRF model was trained on the New England Corpus, a dataset collected by Snow et al. (1996), which contains over 55,000 manually annotated utterances. This dataset was collected as part of a longitudinal study and is the largest child-caregiver interaction dataset. Nikolaus et. al used various baseline models such as majority classifiers, but when comparing performance of all the models on the test dataset, the CRF model had the highest accuracy of 72.33%. For testing, the CRF model was tested on the North American English-language subset of the CHILDES repository. Given its robust performance and alignment with the sequential nature of conversational speech, we selected this model to automatically annotate our dataset with speech act labels.

### Analysis

#### Production

We first restricted the dataset to child speech by selecting utterances labeled CHI and discarding utterances without a speech act label. We calculated the frequency of each speech act for the two groups: late talkers and typically developing children. To calculate the frequency, we first computed the proportion of each speech act within a single transcript. Then, we calculated the mean proportion for all transcripts within a child’s assigned age group. These results were used to compare the use of speech acts between the late talker and typically developing groups.

To compare the frequency of each speech act between typically developing (TD) and late talker (LT) groups, we computed the proportion of utterances containing each speech act within each transcript and found the average of these proportions per child. These proportions, representing continuous variables between 0 and 1, were then compared between groups using Welch’s two-sample t-test, which accounts for unequal group variances. For each speech act type, we obtained a t-statistic and raw p-value, followed by Benjamini–Hochberg False Discovery Rate correction (α = .05) to adjust for multiple comparisons. Speech acts with adjusted p-values below this threshold were considered significant, and the 20 most significant acts were reported in order of corrected p-value.

Because the analysis compares average speech act frequencies across transcripts, treating proportions as continuous measures, Welch’s t-test is statistically appropriate. The test assumes approximate normality of group means, and this is satisfied here due to the moderate sample size and continuous nature of the proportions. Thus, Welch’s *t*-test provides a valid and interpretable method for identifying group-level differences in speech act usage.

#### Classification of LT and TD Groups

To assess whether the use of speech acts can effectively distinguish between typically developing (TD) children and late talkers (LT), we explored a classification approach using speech act distributions as features. Each sample in our dataset was represented as a record, with features corresponding to the proportions of different speech act types.

We evaluated six standard classification algorithms: logistic regression, support vector machines, k-nearest neighbors, Gaussian naive Bayes, random forest, and XGBoost. The dataset was split into training and test sets using a stratified 80/20 split. Each model was trained on the training data and evaluated on the test set using accuracy, F1 score, precision, and recall as performance metrics. To assess model stability and robustness, we applied stratified 4-fold cross-validation and reported the average scores across folds. We selected four folds to balance evaluation reliability while avoiding overfitting, given the relatively small size of our dataset. All classifiers were run with default hyperparameters.

To improve the performance of the best performing classifier, we performed hyperparameter tuning using randomized search. Hyperparameter tuning involves selecting optimal model settings such as the number and depth of trees, splitting criteria, and feature selection strategy, that are not learned from the data but significantly influence predictive performance. Randomized search offers a more efficient alternative to grid search by sampling a fixed number of parameter combinations from predefined ranges. (Franceschi et al., 2025)

We wanted to explore if adding linguistic information in addition to the speech acts could aid our ability to classify LT and TD groups. We selected three additional linguistic features to include for each sample: phonological neighborhood density (phonology), content word frequency (lexical features), and mean length of utterance (as a measure of sentence-level complexity). The process for computing these features followed a similar approach to how we calculated speech act frequencies, averaging the values across the selected transcripts for the sample. For longitudinal datasets, we randomly sampled from the age groups the child appeared in and averaged the feature values from transcripts in that age group to get a value for these metrics. In addition to these linguistic features, we also included assigned age groups and sex of the child as additional metadata for training the models.

#### Speech act clustering

As an exploratory step, we investigated the potential benefit of reducing redundancy and overlap among speech act features by applying hierarchical clustering. This idea was motivated in part by observations in Nikolaus et al. (2022), where the authors note that “many pairs of speech acts are either very similar or hierarchically related. More concretely, there are pairs of speech act categories that describe overlapping communicative act appears to be covered by the other broader act.” To address this, we applied hierarchical clustering to group similar speech acts together. We began by computing the absolute pairwise correlation matrix between all features, then transformed it into a distance matrix by subtracting each value from one. This transformation allowed us to treat highly correlated features as being closer in a hierarchical sense. We applied complete linkage clustering on the resulting distance matrix and visualized the output with a dendrogram, which revealed several tight clusters among speech acts. In addition, we created dendrograms to compare before and after feature clustering. The clustering allowed us to also compare how the clusters overlap with the manual speech act groupings.

### Comprehension

In addition to exploring how children produce speech acts, we also aimed to assess how they comprehend and respond to them. Prior research has focused heavily on production deficits in late talkers (LT), but we hypothesize that their difficulty may extend to comprehension, specifically, in recognizing and responding with an appropriate speech act when addressed by a caregiver. To investigate this, we implemented a comprehension analysis using adult–child adjacency pairs. We identified instances where an adult utterance was immediately followed by a child utterance within the same transcript, forming potential response pairs. For each pair, we extracted the adult’s speech act and the subsequent child response, and checked whether the response is considered an appropriate match using a predefined contingency mapping derived from prior work of Nikolaus et al. If the child’s speech act aligned with the expected contingent response, it was flagged as contingent. To quantify comprehension at the individual level, we calculated the frequency a child produces a contingent response for each response pair.

## Results

### Production

Figure 1 lists the 55 speech acts detected by the CRF model across the datasets and their respective frequency. Aside from ‘YY’ (word-like utterance without clear function), the top five most commonly used speech acts by both groups were ‘ST’ (make a declarative statement), ‘SA’ (answer a *wh*-question with a statement), ‘RP’ (request an action for hearer), ‘AA’ (answer yes/no question) and ‘QN’ (ask a *wh*-question).The top three speech acts: ST, YY and SA made up over half of the utterances spoken by the children across the datasets.

**Figure 1.**
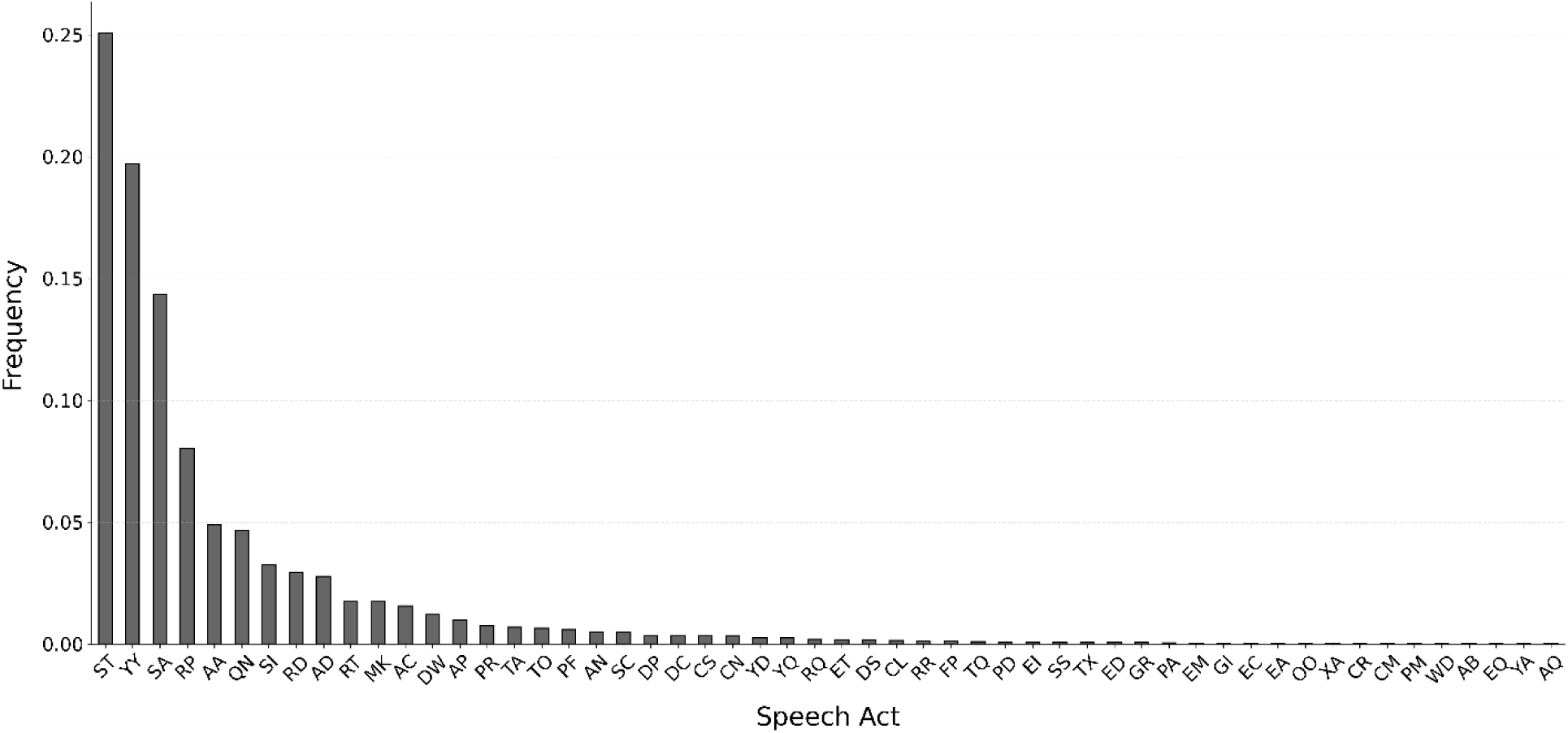
Distribution of Frequencies of Speech Acts Across the Selected Datasets.

Figure 2 illustrated the distributions of the number of distinct speech acts used by the children of both groups across the four age groups. The dashed line indicates the median. We saw that at initial stages of development, the number of distinct speech acts used was centered around 21, but as the age increased, the median decreased to 13, suggesting that as children get older, they rely on less diversified speech acts to achieve their communicative purposes when interacting with other people.

**Figure 2.**
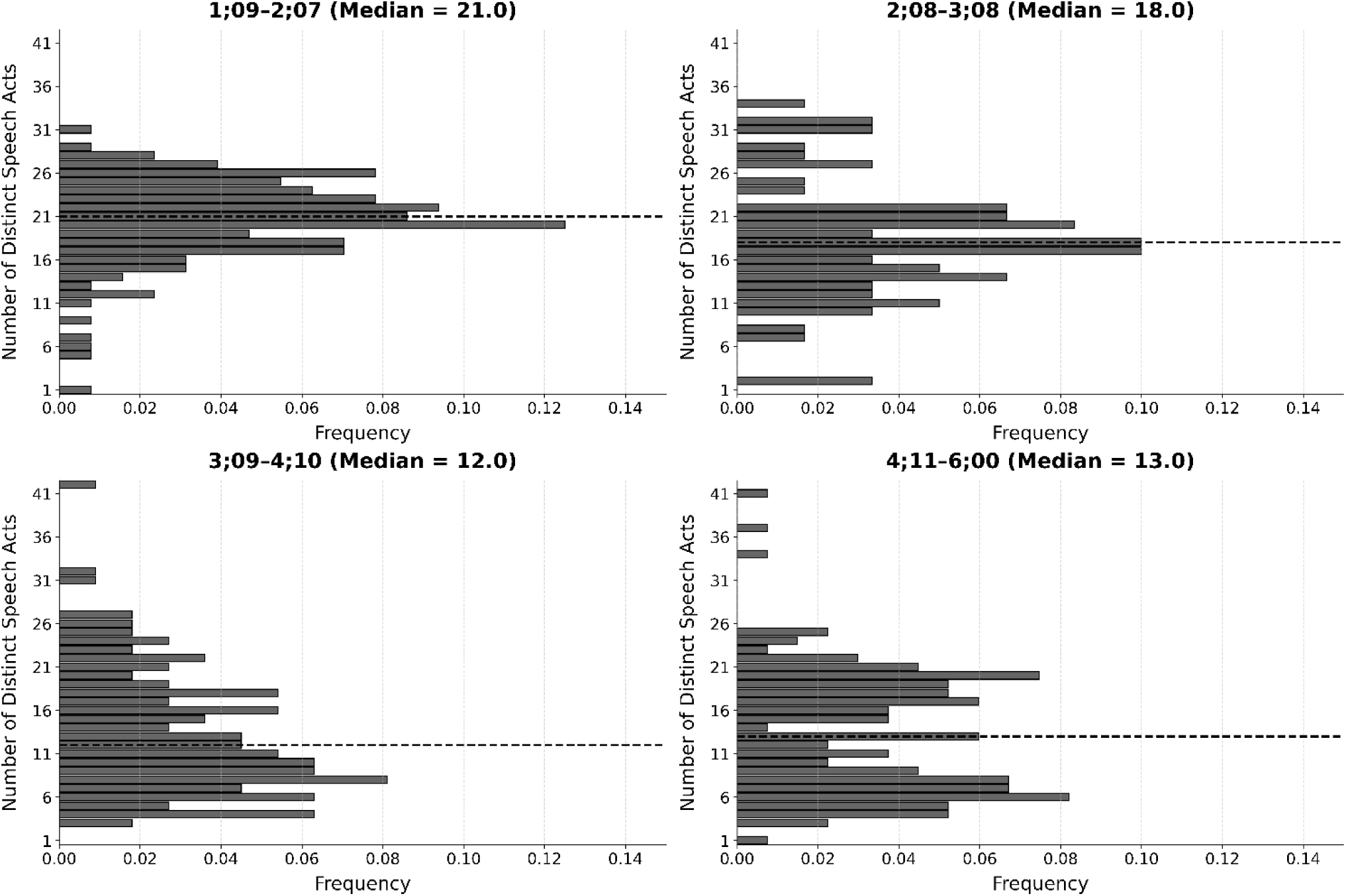
Frequency of distinct speech acts across age groups.

To compare how speech acts vary across age groups and between TD and LT groups, we listed the proportions of each speech act in Figure 3. A common feature of the graph is the distribution of frequencies is concentrated in certain speech acts while the other speech acts occur much less frequently. More specifically, the LT group had a higher proportion of using unclear utterances (YY) than the TD group across all four age groups, but both groups used fewer ‘YY’s when growing older. In contrast, the ‘ST’ showed an opposite trend. The TD group used more declarative statements than the LT group, even starting from age 1;09, and both groups tended to use more as they grew older. The TD group asked the *wh*-questions (QN) and answered yes/no questions (AA) more often to maintain interactions in the first two age groups, but we see this observation switch after age 3;09. TD group was observed making more requests (RP) than LT group in the age group 1;09–2;08, but not for other age groups. We see the reverse in older age groups starting from 2;08. For ‘SA’, we see that its usage by the TD groups drops significantly after 3;09 compared to the LT group.

**Figure 3.**
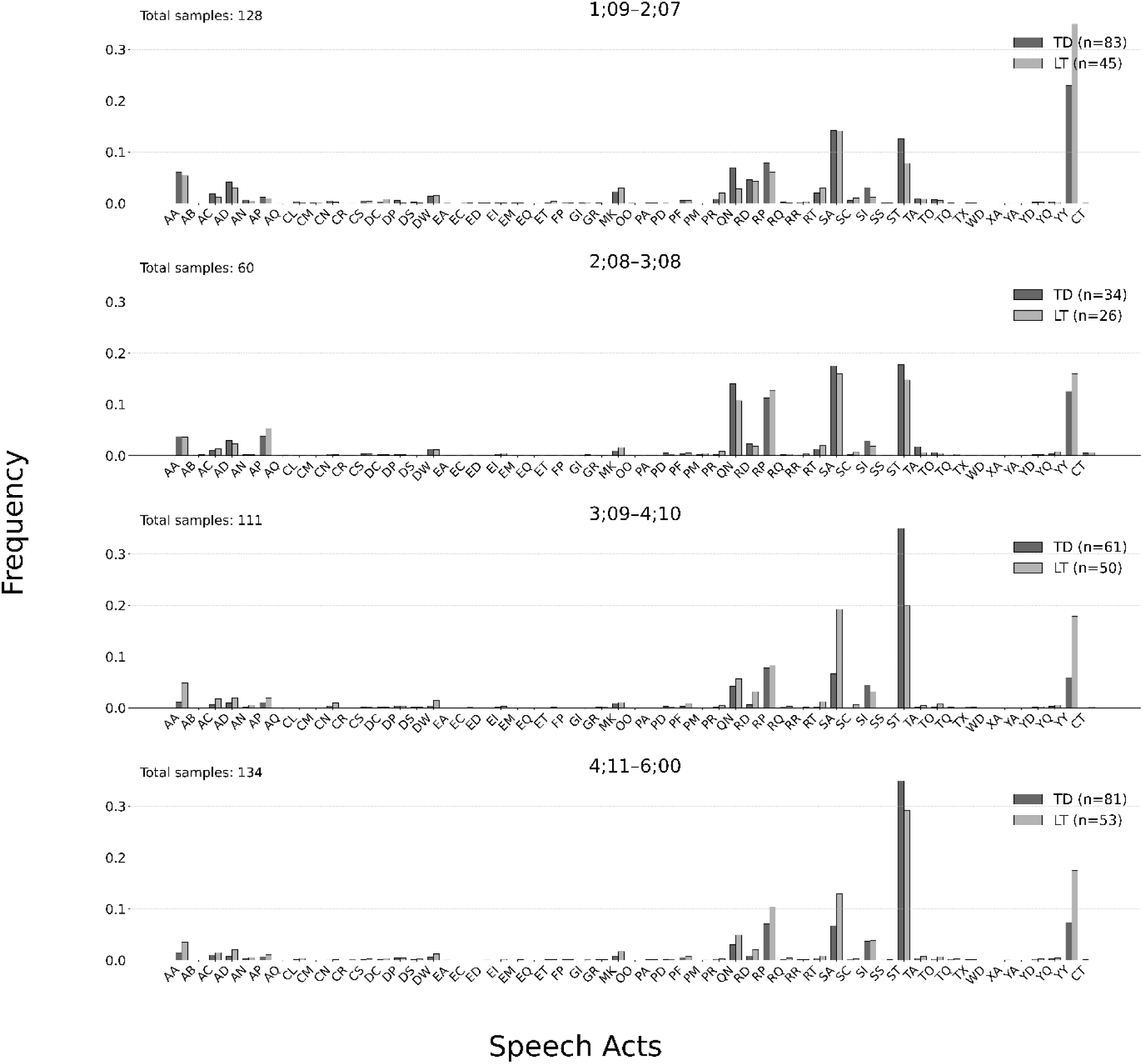
Frequency distribution of speech acts between age groups, and TD/LT groups.

Table 2 reported the t scores and *p*-values of the two-sample t-tests with multiple comparison corrections on the frequency of speech acts between groups and divided by age group. Negative t-statistics indicate that the speech act is, on average, more frequent in LT than in TD. There was no significant difference in the frequency of speech acts between TD and LT in the age group 2;08–3;08. The overall finding was that the TD children consistently used more ‘ST’ than the LT groups across the three age groups. They also asked more *wh*-questions (QN), expressed more intents to act out (SI), ask more permissions to carry out acts (FP) and declared more make-believe (DP) as young as 1;09–2;07 and declared more make-believe as early as 1;09 to 2;07. The LT children, on the other hand, did not begin to use varied speech acts to regulate interaction and exchange information until after 3;09. They also used more word-like utterances without clear function (YY) throughout all age groups than the TD children.

**Table 2.**
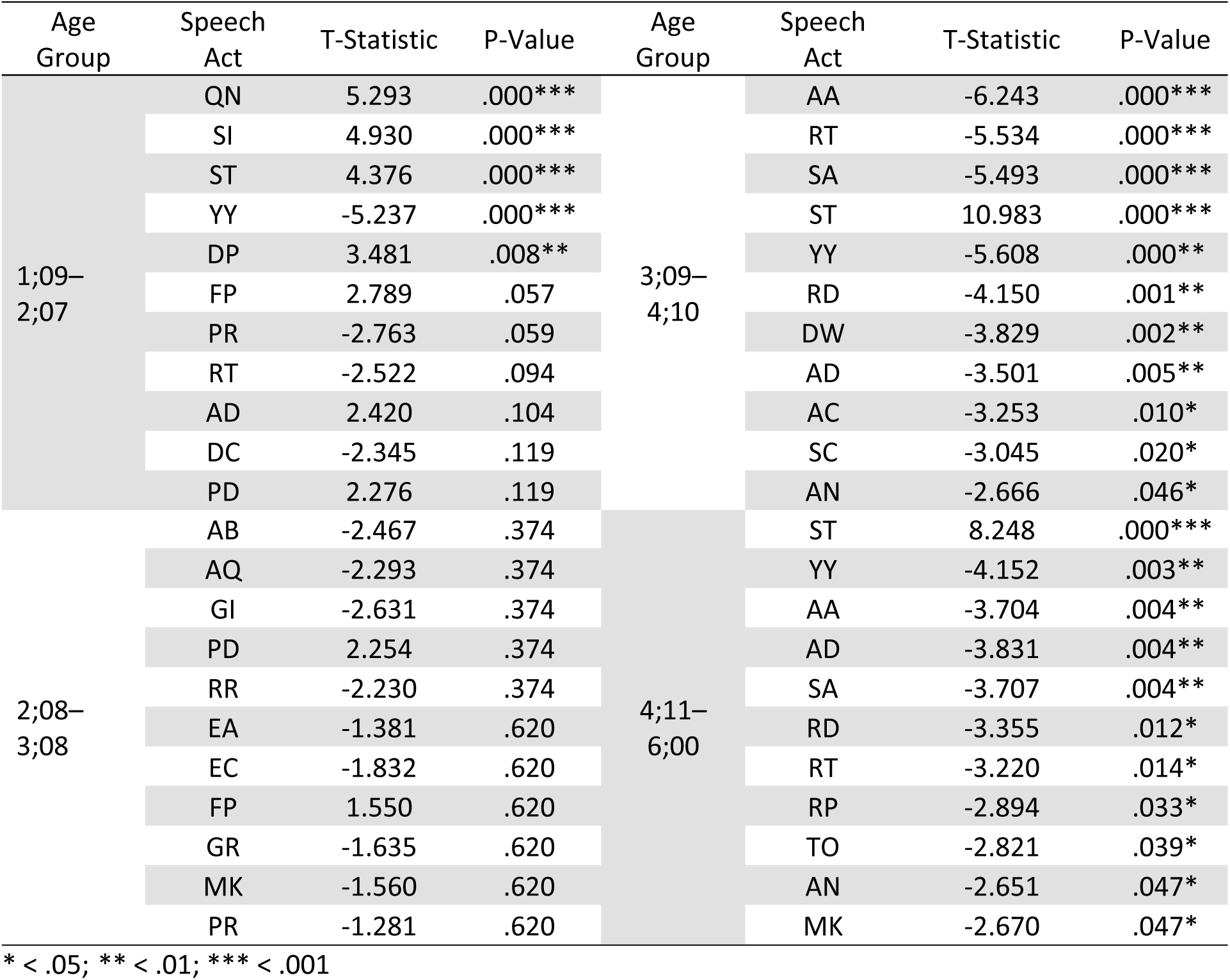
Comparison of speech acts across TD and LT groups.

#### Classification of LT and TD groups

Table 3 listed the accuracy, F1 score, precision and recall as metrics to measure the performance of the five machine learning models we tested. Overall, the Random Forest classifier achieved the highest performance on the four metrics, with an accuracy of .721, an F1 score of .624, precision of .687, and recall of .586. Performance of other models were mediocre. Naive Bayes yielded the lowest performance across all metrics, achieving .642 accuracy and .466 F1, reflecting its limitations in capturing the complex distributions of speech act features. These results suggest that ensemble tree-based approaches, particularly Random Forests, are most effective for leveraging speech act distributions to classify children’s developmental status.

**Table 3.** Results from 4-Fold Cross Validation using speech act features.

| Model | Accuracy | F1 Score | Precision | Recall |
| --- | --- | --- | --- | --- |
| Random Forest | .721 | .624 | .687 | .586 |
| SVM | .700 | .592 | .655 | .546 |
| Logistic Regression | .700 | .593 | .651 | .557 |
| KNN | .695 | .600 | .638 | .575 |
| XGBoost | .677 | .573 | .613 | .541 |
| Naive Bayes | .642 | .466 | .562 | .412 |

We performed hyperparameter tuning on the best model, random forest, to assess how much further we could improve the results. The best-performing configuration used 700 decision trees, a maximum depth of fifteen, log2-based feature sampling, and required a minimum of two samples to split and two sample per leaf, with bootstrap aggregation enabled. We applied 4-fold stratified cross-validations and achieved accuracy of .723, F1 score of .618, precision of .715, and recall of .558. These results show that tuning improved accuracy, F1, and precision, with a modest gain in recall, enhancing the model’s ability to reliably classify the TD and LT groups.

We further explored the contributions of linguistic features (phonological neighborhood density, content word frequency, mean length of utterance), and demographic features – age and sex to model classification in conjunction with the speech acts. We observed improved performance for several models, among which XGBoost achieved the highest overall accuracy at .738, with an F1 score of .663, precision of .689, and recall of .642, indicating balanced and consistent performance. KNN achieved the second highest overall accuracy (.722, F1 = .640), while SVM produced the highest recall (.839) but with the lowest accuracy (.583).

These findings suggest that incorporating linguistic features, age, and sex provides a measurable performance boost, particularly for ensemble and instance-based models like XGBoost and KNN. Ensemble models like XGBoost and instance-based models like KNN improved with the inclusion of age, sex, and linguistic features possibly because these methods capture more complex patterns than speech acts alone. XGBoost can model nonlinear interactions, such as how the same number of speech acts may indicate different outcomes depending on a child’s age, sex, or linguistic features (Chen & Guestrin, 2016). KNN may benefit from a richer feature space that includes age, sex and linguistic features, allowing it to group children more accurately with peers who share similar developmental and linguistic profiles (Cover & Hart, 1967). By contrast, simpler models such as Logistic Regression or Naive Bayes are less able to account for these interactions, which may explain why their performance gains were more limited.

Given that XGBoost was our best-performing model using this expanded feature set, we further explored how much its performance could be improved through hyperparameter tuning. Using the same procedure as before, we performed hyperparameter tuning using random search. Tuning led to measurable performance gains: accuracy improved from .738 to .766, F1 score from .663 to .703, precision from .689 to .721, and recall from .642 to .688. The best model used 300 estimators, a learning rate of .2 and maximum tree depth of 10.

#### Hierarchical clustering of speech acts

The dendrogram in figure 4 shows seven cluster-level feature sets. The most similar pair is C3– C7, which merge at the lowest height: both are dominated by Information Exchange, with C3 centered on assertions/answers plus discourse devices (e.g., ST, SI, DW, RT, SC; AA/AD) and C7 emphasizing questioning with brief replies and protests/requests (e.g., QN, YQ, YA, GR, WD, PD, AQ, GI). By contrast, C1 (AN, CL, FP, PA) merges only near the top of the tree, indicating it is least like the others; its narrow mix of answers (AN, PA), directing attention (CL), and justification/request (FP) contains the most dissimilar grouping.

**Figure 4.**
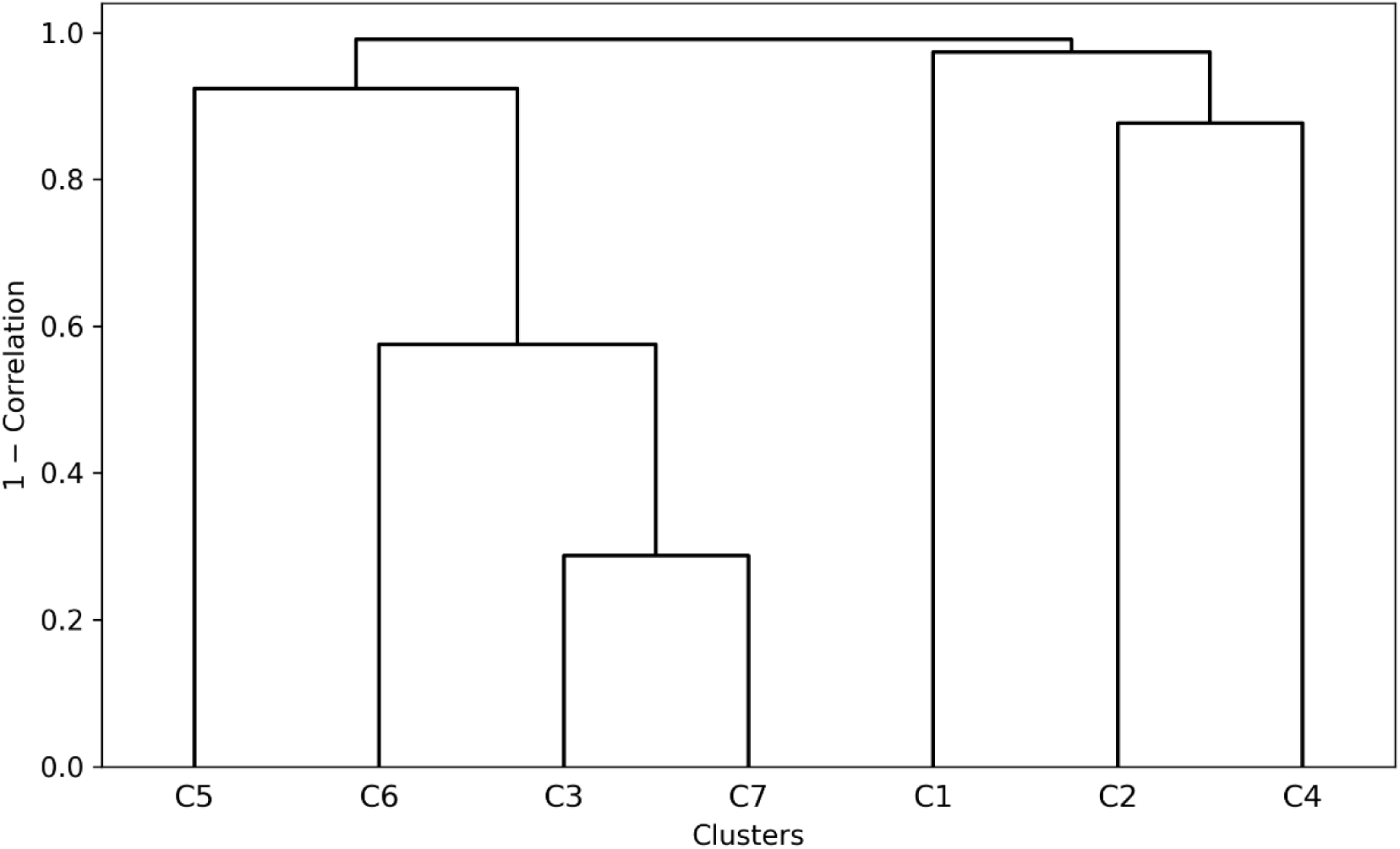
Dendrogram of Hierarchically Clustered Features.

We compared the results of our data-driven hierarchical clustering (C1–C7, Table 4) with the 7 theoretical categories of tier 1 speech acts (Appendix Table 1). One cluster is close to a declarations group (C4: DP, TX with DS), but most clusters are mixed across manual categories—for example, C2 pairs AN (answering) with CL (joint attention), while C3, C6, and C7 blend information-exchange items (asserting, questioning, answering, repairing, evaluative/declarative) with interaction-regulation, discourse/organizational devices, and miscellaneous codes.

**Table 4.** Groupings after hierarchical clustering.

| Cluster | Features |
| --- | --- |
| 1 | AN, CL, FP, PA |
| 2 | CM, CR, DC, DP, DS, OO, PR, RR, TQ, TX, YD |
| 3 | AA, AC, AD, DW, MK, PF, RD, RT, SC, SI, ST, TO, YY |
| 4 | CS, EA, EC, ED, EM, ET, SS |
| 5 | CN, EQ, SA, TA, XA |
| 6 | AB, AP, CT, EI, PM, RP, RQ |
| 7 | AQ, GI, GR, PD, QN, WD, YA, YQ |

The meanings of the speech act abbreviations can be understood from the work of Ninio et.al (1994).

We used these clusters to perform the same classification workflow as before, if reducing overlapping speech acts would improve performance. After reducing the feature space into seven clusters, overall model performance declined: Random Forest dropped from .721 to .624 accuracy, SVM from .700 to .640, and Logistic Regression from .700 to .621. XGBoost, Decision Tree, and KNN showed similar decreases, while Naive Bayes remained largely unchanged. These results suggest that clustering sacrifices predictive resolution, indicating that the finer-grained INCA-A speech act categories provide stronger discriminative power than broader theoretical groupings.

### Comprehension

We performed four independent sample t-tests with multiple comparison corrections to compare the proportions of contingent speech acts produced by the LT and TD children. There was a significant group difference in the youngest age group (1;09–2;07) (Figure 5), with TD children showing higher contingency rates than their LT peers (t = 3.249, corrected *p* = .002).

**Figure 5.**
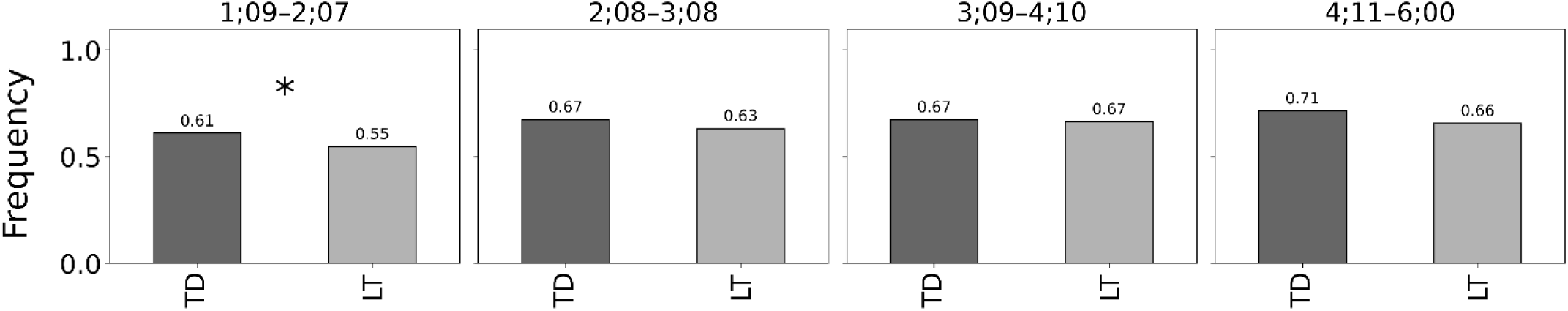
Contingent Response frequency across age groups.

## Discussion

This study investigates how speech acts differentiate typically developing and late-talking preschoolers using automated INCA-A labeling based on the CRF model and machine learning across nine CHILDES corpora comprising 433 children between 1;09 and 6;00. We applied age-stratified sampling, tested production differences with Welch’s t-tests and FDR, indexed comprehension via adult–child adjacency pairs, and built cross-validated classifiers with model interpretation. The TD children produced more declarative statements and *wh*-questions and showed higher rates of contingent responses; the LT children produced more unclear or word-like utterances and were less competent in comprehending speech acts at 1;09–2;07. Reliable classification of the LT and TD children depended on configurations of co-occurring acts rather than single-act frequencies.

Two salient differences between the LT and TD children were robustly detected in the 1;09– 2;07, 2;08–3;08, 3;09–4;10, and 4;11–6;00 age groups across a variety of contexts for adult-child interactions, ranging from naturalistic toy play, free play with game, storytelling from pictures to more controlled elicited picture description task. First, the LTs produced more word-like utterances without clear function (YY) than the age-matched the TDs regardless of the age and started to rely more on the speech acts that can regulate interaction (AA, AD, SA, RD, RP, TO) and exchange information (AA, SA, DW) until after age 3;09. Our results did not support the earlier report that the repertoire of communicative acts in TD increased with age (Bergey et al., 2024). Instead, we found that preschool LT children can resort to multiple types of speech act tools to compensate for the deficient information density in their speech production. They made more responses to directives/assertion, more requests and showed more disagreements in their interactions. On the other hand, the TD children consistently produced more declarative statements (ST) across the three age groups. As ‘ST’ has the dual functions of co-referencing and asserting, it implies that the TD children are more capable of directing hearers’ attention to a shared context and use joint attention to create a basis for mutual understanding to facilitate communication. They also preferred to ask more *wh*-questions (QN), expressed more intents to act out (SI) and declared more make-believes which is a representation of theory of mind as young as ages 1;09 to 2;07. These contrastive differences show that the early LT *can* use varied speech acts to achieve their communicative goals, but they tend to be more ‘speaker-centered’ when resolving communication needs, while the TD children are more ‘hearer-centered’ by shared focus to engage the hearer and maintain cooperative communicative process.

Our study also showed that speech act played a dominant role in predicting the LT status at the preschool age with the best accuracy of 72.1% in the Random Forest model. This model can be generalized well on the test data with accuracy of 74.2%. The finding reinforces the relevance of using speech act markers to flag early intervention for preschool LT children. Our feature engineering by including the phonological, lexical, grammatical, and demographic factors like age and sex produced a slight improvement of accuracy for the XGBoost (73.8%) and KNN (72.2%) models. The improvement revealed more complex and nonlinear interactions among linguistic, demographic and pragmatic features. The top five predictive features contributing more than 5% to the model splitting power (i.e. ST, YY, SA, QN and AD) were also ranked highly in the overall proportion of production in utterances across the two groups (Figure 1).

Lastly, although the TD and LT children showed differences in producing speech acts across all age groups, only TD children in the youngest age group showed higher contingency rates than their LT peers in comprehension. The result indicates that comprehension differences may emerge early in development (Snow et al., 1996; Nikolaus et al., 2022), supporting the idea of a critical window of speech development when deficits in the receptive knowledge of speech act functions may influence the full development of production skills. As our evidence suggested, the window could start as early as the age of 1;09. The lack of differences in the other three age groups is possibly related to the greater competence of the LT children in producing varied speech acts, which in turn helps them to obtain more speech cues to improve understanding. Our result implies that drilling linguistic competency like phonological, lexical and grammatical knowledge may yield less optimal therapeutic outcomes at this early stage as these linguistic skills develop much later than speech act knowledge (Stephens & Matthews 2014). Early intervention of the receptive language ability of the LT children needs to prioritize helping these children recognize the distinct interactional functions of speech acts through multimodal methods like facial expressions, body language, paralinguistic and contextual cues.

Another aim of our study was to further examine the relationship among speech acts of the INCA-A scheme by comparing the theoretical classification (Table A1) to the data-driven clusters. The assumption came from Nikolaus et al. (2022) that reducing redundancy and ambiguity in speech act categorization should better characterize the differences between the LT and TD children in terms of model performance. However, our results failed to support this assumption due to decreased accuracy in classification. Moreover, there is no clear mapping between the 7 clusters generated by our data and theoretical taxonomy. Each tier 1 pragmatic function was mixed with multiple clusters of speech acts. Notably, cluster 3 integrates Information Exchange with discourse organization and joint attention, with smaller contributions from social ritual and miscellaneous acts, whereas cluster 7 is dominated by questioning within Information Exchange alongside interaction-regulation responses/protests. All 7 clusters contributed to interaction regulation and information exchange. One plausible explanation for the mismatch is that the two clustering approaches rely on different organization principles. Theoretical taxonomy uses the speakers’ intended illocutionary force and interactive function, while the data-driven approach derived the clusters based on the correlations (i.e. similarity) calculated from the average proportion of utterances across transcripts per child. High correlation may indicate a high co-occurrence rate in speech samples. For example, FP (ask for permission to carry out act) and PA (permit hearer to perform act) were grouped into cluster 1, suggesting that the two speech acts were often used in tandem by the children to construe the utterance’s illocutionary force and enact the perlocutionary force on hearers. This discrepancy of the two frameworks further points out the necessity of adapting the feature descriptors from a multidimensional perspective, instead of simply relying on usage frequency to represent speech act function.

## Conclusion

While the number of studies on LT children is picking up in recent years, utilization of large-scale LT corpora combined with large language models to tap into the communicative deficits of the preschool LT children can help medical professionals to identify potential markers for early diagnosis and intervention of speech delays in infants and toddlers. Investigation of speech acts that serve communicative functions among the LT children can yield valuable insights into the role of social interactions in influencing the developmental trajectories of language and cognition of these children. A deeper understanding of these effects will drive progress in speech therapy to provide individualized treatment protocols for children who are late bloomers and who may experience persistent language delay at school age.

## Data, Code and Availability Statement

Source code of all the analyses done, figures and outputs of classification models are publicly available at: https://github.com/girwandhakal/Speech-Act-Analysis

Datasets used for the analyses are available at https://talkbank.org/childes/ and are accessible upon registration.

## Appendix

**Table A1.**
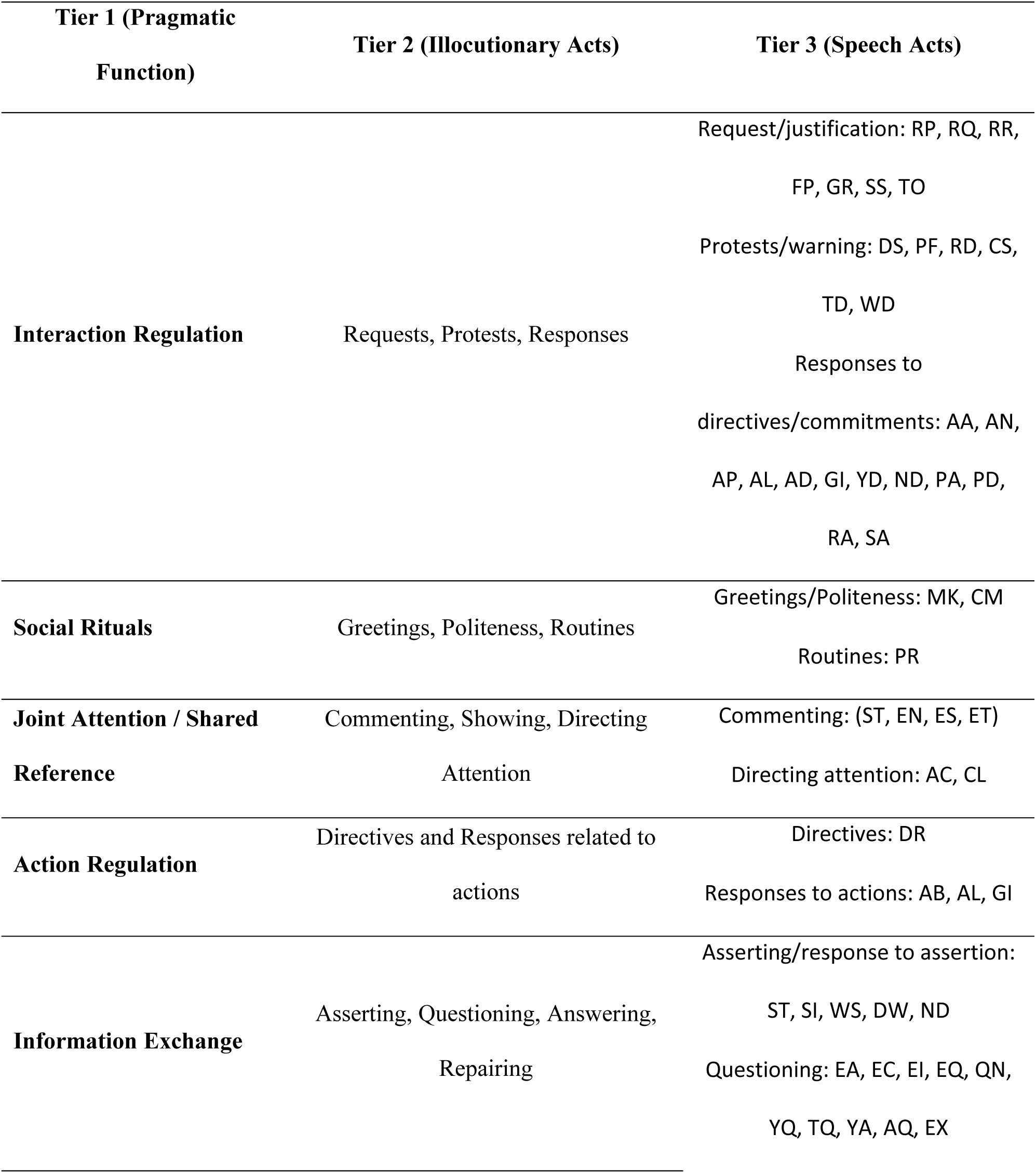
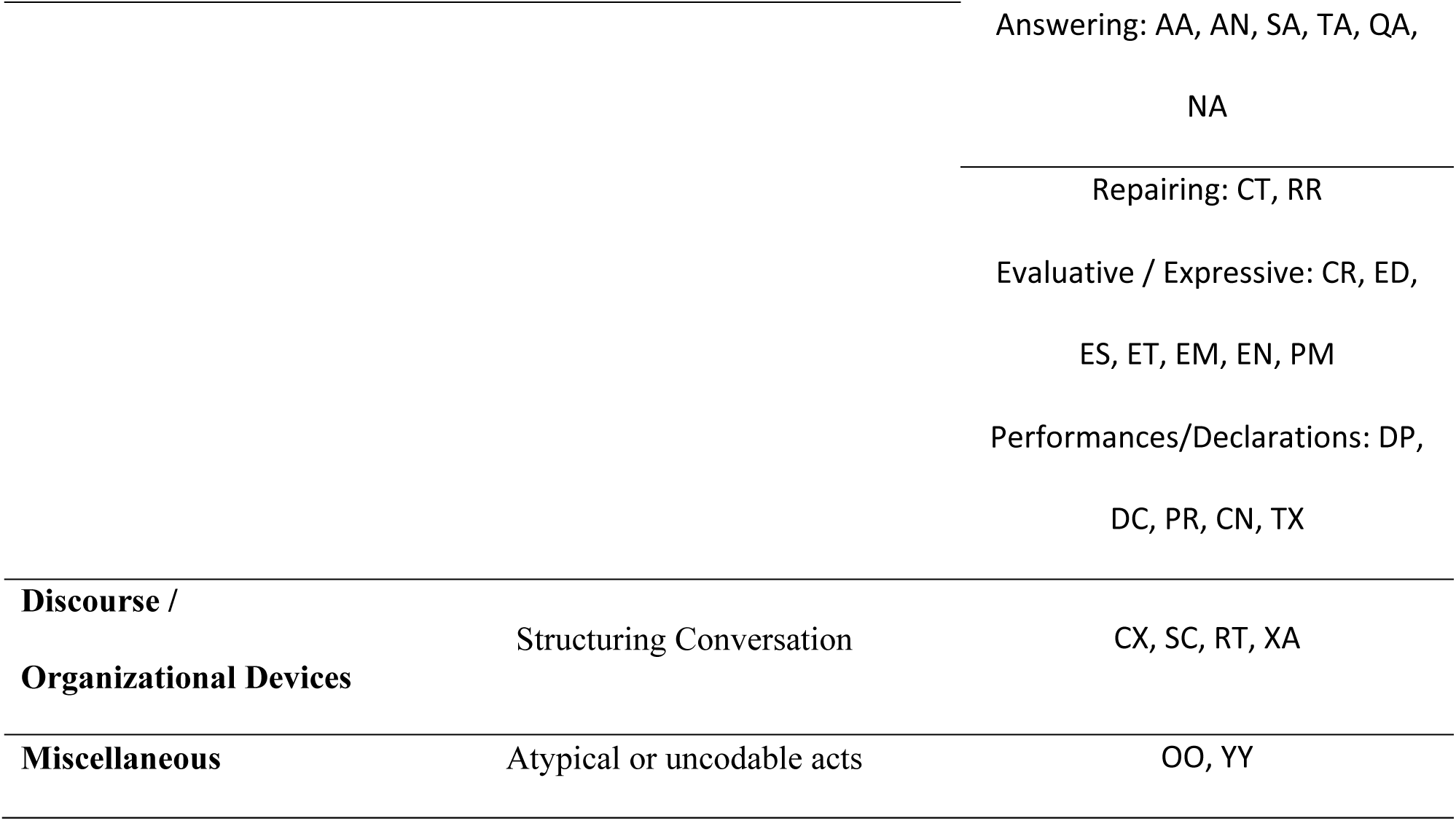
Tier 1–3 Structure of the INCA-A Coding Scheme.

## Notes

### Competing Interest Statement

The authors have declared no competing interest.

